# Deep learning modeling of ribosome profiling reveals regulatory underpinnings of translatome and interprets disease variants

**DOI:** 10.1101/2024.02.26.582217

**Authors:** Jialin He, Lei Xiong, Shaohui Shi, Chengyu Li, Kexuan Chen, Qianchen Fang, Jiuhong Nan, Ke Ding, Jingyun Li, Yuanhui Mao, Carles A. Boix, Xinyang Hu, Manolis Kellis, Xushen Xiong

**Affiliations:** The Second Affiliated Hospital, Zhejiang University School of Medicine, Hangzhou 311121, China; Liangzhu Laboratory, School of Medicine, Zhejiang University, Hangzhou 311121, China; State Key Laboratory of Transvascular Implantation Devices, The Second Affiliated Hospital, Zhejiang University School of Medicine, Hangzhou 311121, China; Computer Science and Artificial Intelligence Lab, Massachusetts Institute of Technology, Cambridge, MA 02139, USA; Sir Run Run Shaw Hospital, Zhejiang University School of Medicine, Hangzhou 310016, China; Department of Biomedical Informatics, Harvard Medical School, Boston, MA 02115, USA

## Abstract

Gene expression involves transcription and translation. Despite large datasets and increasingly powerful methods devoted to calculating genetic variants’ effects on transcription, discrepancy between mRNA and protein levels hinders the systematic interpretation of the regulatory effects of disease-associated variants. Accurate models of the sequence determinants of translation are needed to close this gap and to interpret disease-associated variants that act on translation. Here, we present Translatomer, a multimodal transformer framework that predicts cell-type-specific translation from mRNA expression and gene sequence. We train Translatomer on 33 tissues and cell lines, and show that the inclusion of sequence substantially improves the prediction of ribosome profiling signal, indicating that Translatomer captures sequence-dependent translational regulatory information. Translatomer achieves accuracies of 0.72 to 0.80 for *de novo* prediction of cell-type-specific ribosome profiling. We develop an *in silico* mutagenesis tool to estimate mutational effects on translation and demonstrate that variants associated with translation regulation are evolutionarily constrained, both within the human population and across species. Notably, we identify cell-type-specific translational regulatory mechanisms independent of eQTLs for 3,041 non-coding and synonymous variants associated with complex diseases, including Alzheimer’s disease, schizophrenia, and congenital heart disease. Translatomer accurately models the genetic underpinnings of translation, bridging the gap between mRNA and protein levels, and providing valuable mechanistic insights toward mapping “missing regulation” in disease genetics.

## Introduction

Gene expression is a complex process that involves transcription and translation. Although message RNA (mRNA) level is predictive of the corresponding protein level, the discrepancy between these two expression layers has increasingly been realized^1–4^. Studies have revealed that the correlation between mRNA and protein abundance is roughly 0.6, as represented by Pearson correlation coefficient. The transcription-translation discrepancy varies across different cell types and tissues, with a range of correlation between 0.39 to 0.79 in human^3,5^. However, the mechanisms and regulatory rules underlying such context-dependent discrepancy remain largely uncharted.

Over 93% of human disease-related variants do not alter protein sequences^6,7^. Enormous efforts have been made to understand the mechanisms of how these non-protein-coding variants contribute to human traits and phenotypes, with the majority of the studies focusing on interpreting disease loci at the layer of mRNA expression^8–10^. However, a large fraction of non-coding variants remain uninterpreted, leading to the puzzle of “missing regulation” in the field of disease genetics^3,11^. Given the discrepancy between mRNA and protein, using mRNA level to represent the functional output of a gene may lead to the neglect of biological mechanisms at the translation level. However, there are relatively limited studies focused on the level of translation, although protein is arguably more closely related to phenotype^12,13,14,15^. In fact, a considerable fraction of disease-relevant variants (∼58% based on the GWAS catalog) are located in gene body instead of promoter or enhancer and likely exert little effect on transcription. The regulatory mechanisms of the variants located in the 5’ untranslated region (5’ UTR), 3’ untranslated region (3’ UTR) and synonymous mutations are largely unclear.

Ribosome profiling is a high-throughput technique that monitors the footprints of ribosome movement on mRNA, which is one of the most well-established proxies for protein synthesis^16,17^. There have been studies utilizing ribosome profiling to build connections between genetic variants and translation, namely ribosome profiling QTLs and translation efficiency QTLs^18,19^. However, the identified genetically-driven translational regulation loci are very limited thus far, likely due to the technical hurdles and the high cost of ribosome profiling experiments^18,19^. By contrast, RNA sequencing (RNA-seq) is more cost-effective and easier to handle experimentally. Therefore, building a model that predicts ribosome profiling from RNA-seq signal will provide an opportunity to interrogate the translation process even with transcription information available only, and meanwhile help understand the regulation underlying the gap between transcription and translation.

The advent of deep learning has largely facilitated the prediction of various layers of molecular phenotypes and the investigation of the regulatory information underlying genomic sequence^20–23^. Likewise, several deep learning models have been developed to predict ribosome profiling density along mRNA molecules using gene sequence as an input^24–26^. However, while there has been a variety of known and presumably more unknown sequence elements in 5’ UTR and 3’ UTR that regulate the translation process, the existing models are primarily focused on coding sequence, with the untranslated regions largely neglected. Transformer applies attention modules and is frequently used in natural language processing (NLP) and computer vision to capture long-distance relationships^27^. Given that mRNA can be translated in a “looping form” and that tertiary structure of mRNA may affect translation^28–30^, the attention mechanism holds the promise for identifying distal interactions within an mRNA molecule during translation. Most importantly, none of the currently available methods for ribosome profiling prediction integrate transcriptome information as input, therefore losing the ability to study the relationship between transcription and translation.

Here, we present Translatomer (a portmanteau of translatome and transformer), a transformer-based multimodal deep learning framework that predicts ribosome profiling track using gene sequence and cell-type-specific RNA-seq as inputs (**Fig. 1a-b**). We curate 33 ribosome profiling and the paired RNA-seq across a wide range of tissues and cell types for model training, which enables the reservation of tissue/cell-type specificity of translation from the corresponding mRNA information. Translatomer outperforms the state-of-the-art deep learning models in both the prediction accuracy and the ability to capture translation regulatory information. We show that the prediction accuracy is increased by 5.3% upon the inclusion of sequence as an additional input along with RNA-seq. Translatomer is capable of accurately predicting the ribosome profiling signal from unseen cell types, indicating the generalizability of the model. To enable the interrogation of the effect of genetic variants on translation, we develop an *in silico* mutagenesis approach that estimates the translational alteration upon sequence mutation. We demonstrate that the untranslated regions (UTRs), which do not directly affect protein sequences, contribute to evolutionary constraints by affecting the translation efficiency. We then utilize Translatomer to interpret the mechanism of the synonymous and the UTR-located variants curated by ClinVar that are reportedly relevant to human diseases. Intriguingly, we characterize 3,041 genetic variants that are related to brain diseases such as Alzheimer’s disease, amyotrophic lateral sclerosis, schizophrenia, and autism, as well as heart diseases including cardiomyopathy, congenital heart disease, and heart failure.

**Figure 1.**
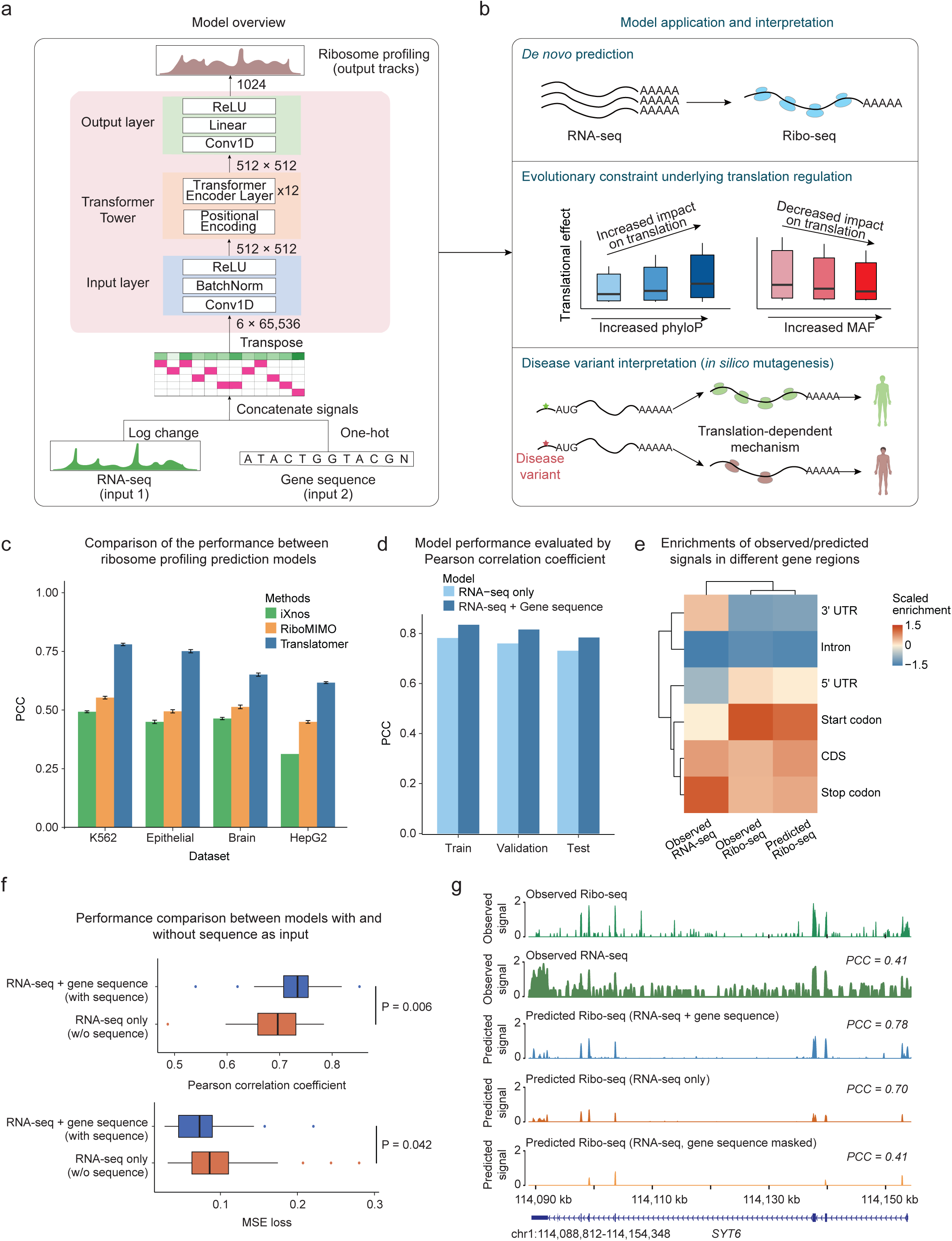
Sequence information bridges the gap between transcription and translation. a) Overview of the multimodal and multi-input design of Translatomer. The model takes in RNA-seq and gene sequence as inputs to predict ribosome profiling signals. b) The applications of the model, including *de novo* prediction for unseen data, evolutionary constraint interrogation and translation-dependent disease variant interpretation. c) Comparison of the accuracy between Translatomer and other cutting-edge ribosome profiling prediction models, including iXnos and RiboMIMO, using an 11-fold cross-validation strategy in K562, epithelial cells, brain and HepG2 datasets. Translatomer consistently outperforms the other models across different cell type contexts. d) The training, validation and testing Pearson correlation coefficients of the RNA-seq only model and the full model across the 33 cell lines and tissues. e) Enrichments of the observed RNA-seq, and the observed and predicted ribosome profiling signals in different gene regions. Hierarchical clustering reveals that the predicted ribosome profiling resembled the observed ribosome profiling signal. f) Pearson correlation (upper) and MSE loss (bottom) between the observed versus the predicted ribosome profiling across the 33 datasets for the full model with sequence information and the RNA-seq only model. Wilcoxon rank-sum test is used for the comparison between the two models. g) Example tracks showing the observed/predicted ribosome profiling and RNA-seq signals in the *SYT6* gene. Pearson correlation coefficient against the observed Ribo-seq signal is labeled at the top right corner for each model.

## Results

### The multimodal design of Translatomer

To bridge the gap between transcriptional and translational levels, we developed Translatomer, a transformer-based deep learning framework that integrates gene sequence with RNA-seq data as inputs to model translation and to understand the regulatory elements in sequence during ribosome movement (**Fig. 1a-b**). Translatomer is composed of two input encoders that take in RNA-seq and gene sequence, a one-dimension (1D) convolution block (Input layer), a transformer tower consisting of a 12-layer self-attention block and a positional encoding layer, and an output layer (**Fig. 1a, Extended Data Fig. 1a, Methods**). The input layer tokenizes the concatenated input of RNA-seq and one-hot-represented gene sequence into 512 tokens, which were then fed into the transformer tower to learn the distal interaction between translational regulatory elements. Lastly, we decode the output by the transformer tower and predict ribosome profiling coverage (**Fig. 1a, Methods)**.

We first compared Translatomer with the state-of-the-art models for ribosome profiling prediction, including a neural network regression model iXnos^26^ and a gated recurrent unit (GRU)-based network RiboMIMO^31^. We selected the ribosome profiling signals from four distinct cell lines and tissues, including K562, epithelial cells, brain, and HepG2, and carried out extensive benchmarking using an 11-fold cross-validation (**Methods**). While we observed accuracies ranging from 0.31 to 0.49 for iXnos, and 0.45 to 0.55 for RiboMIMO across different datasets, Translatomer shows substantially increased accuracies ranging from 0.62 to 0.78 (**Fig. 1c**). The substantial increase of prediction accuracy is likely ascribed to multiple aspects, including the employment of transformer layers in the model, the inclusion of untranslated regions (UTRs) sequence in addition to coding regions, and the inclusion of RNA-seq signal (**Extended Data Fig. 1b**). Given the essential roles of 5’ UTR and 3’ UTR in regulating translation, the addition of these UTR sequences and the employment of transformers together enable the model to utilize and to capture the embedded regulatory information in these regions. The inclusion of RNA-seq signal provides a cell-type/tissue context for accurate translation modeling, and more importantly, provides a unique opportunity to study the sequence-dependent relationship between transcription and translation. These optimizations together greatly enhance the ability of Translatomer in predicting ribosome profiling and in bridging the regulatory gap between transcription and translation (**Extended Data Fig. 1b**).

### Translatomer utilizes sequence information to promote prediction of translatome from transcriptome

We integrated 33 paired RNA-seq and ribosome profiling datasets to train the model, which achieves an accuracy of 0.784, as represented by the Pearson correlation coefficient between the observed and the predicted ribosome density for the held-out chromosome upon the convergence of mean squared error (MSE) loss (**Fig. 1d, Extended Data Fig. 1c-e, Methods**). The predicted ribosome profiling signal is consistent with the observed ribosome profiling, but is distinct from the RNA-seq signal, with strong enrichments at the start codon and negligible signals in the 3’ UTR and intron regions (**Fig. 1e**). This suggests that the model utilizes the sequence information to learn the difference between RNA-seq and ribosome profiling. To quantify the contribution of sequence in modeling ribosome profiling, we trained a model with the same architecture that utilizes only RNA-seq data as input by masking the sequence as zeros for predicting ribosome profiling signals (RNA-seq-only model). This RNA-seq-only model resulted in a Pearson correlation coefficient of 0.731, indicating that the inclusion of sequence information leads to a ∼5.3% (from 0.731 to 0.784) increased accuracy for predicting translation (**Fig. 1d, Extended Data Fig. 1d**). Consistently, the MSE loss shows a 20% reduction from 0.060 to 0.048 (**Extended Data Fig. 1e**). To further evaluate the contribution of sequence information in modeling translation, we applied the model to each of the 33 datasets used for training, with sequence included and excluded as an input, respectively. We observed an elevation of median accuracy from 0.696 to 0.734, and a decrease of median MSE loss from 0.086 to 0.073 upon the inclusion of sequence as an input along with RNA-seq (**Fig. 1f**).

To intuitively demonstrate the importance of sequence information in ensuring the precise prediction of ribosome density, we show the *SYT6* gene as an example, which encodes a synaptotagmin that plays a key role in the acrosomal exocytosis process^32,33^. The Pearson correlation coefficient between the observed RNA-seq and Ribo-seq is 0.41 for *SYT6*, indicating a considerable gap between the two layers (**Fig. 1g**). The full model with both RNA-seq and gene sequence as inputs gave rise to a Pearson correlation coefficient of 0.78 in predicting *SYT6* translation in neurons, while the performance dropped to 0.70 (∼10.3% decrease) when using the model without sequence information (**Fig. 1g**). Additionally, we utilized a second metric that masks the gene sequence in the multi-input model, to evaluate the contribution of sequence information, and again show that the performance is largely attenuated (accuracy = 0.41; **Fig. 1g**). Collectively, the multi-input design of Translatomer extracts sequence information to enhance the accuracy of modeling translation, and to mitigate the gap between transcription and translation by harnessing the regulatory information embedded in the sequence.

### Translatomer enables *de novo* predictions of ribosome profiling

We envision that Translatomer is capable of *de novo* prediction of ribosome profiling for unseen tissues or cell types. This is because our model is trained on 33 datasets across a variety of tissues and cell lines, which allows the model to learn the context-specific regulation from the interaction between gene sequence and RNA-seq signals.

We first applied the model to four human tissues/cell-lines that have not been seen by the model, including prostate, epithelial, hTERT-RPE1, and rhabdomyosarcoma (**Supplementary Table 2**). We fed the model with gene sequence and cell-type/tissue-specific RNA-seq data as inputs to predict the corresponding ribosome profiling signals. While the model without sequence shows lower accuracies, the full model achieves an accuracy of 0.72 for prostate, 0.79 for epithelial, 0.79 for hTERT-RPE1 and 0.80 for rhabdomyosarcoma, with MSE losses of 0.16, 0.11, 0.07 and 0.08, respectively (**Fig. 2a**). We then visualized *KLF6* gene, which encodes a Kruppel-like family transcriptional activator and has been reported to induce apoptosis in epithelial cell^34,35^, as an example to demonstrate the ability of Translatomer in capturing cell-type specificity. The predicted ribosome footprints of *KLF6* in epithelial cells and hTERT-RPE1 (a retinal pigment epithelial cell line) are highly consistent with the observed epithelial ribosome profile (PCC = 0.76 and 0.77, respectively; **Fig. 2b**). The predictions based on the other two non-epithelial tissues/cells, including prostate and rhabdomyosarcoma, show relatively decreased correlation with the observed *KLF6* translation in epithelial (PCC = 0.64 and 0.62, respectively; **Fig. 2b**). Notably, the RNA-seq profiles of *KLF6* between the non-epithelial and epithelial contexts are highly consistent (PCC = 0.92, 0.85 and 0.95), indicating that the cell-type specificity of the predicted Ribo-seq is driven by the specificity at the translational level instead of the transcription level (**Fig. 2c**). As a comparison, the predicted Ribo-seq and the observed RNA-seq signals of *ACTB*, which is a housekeeping gene, are both highly consistent between different tissues (ribosome profiling PCC ranges from 0.88 to 0.91, RNA-seq PCC ranges from 0.96 to 0.98; **Extended Data Fig. 2a-b**). Therefore, our model is capable of preserving different degrees of transcription-translation discrepancy for different genes across tissues.

**Figure 2.**
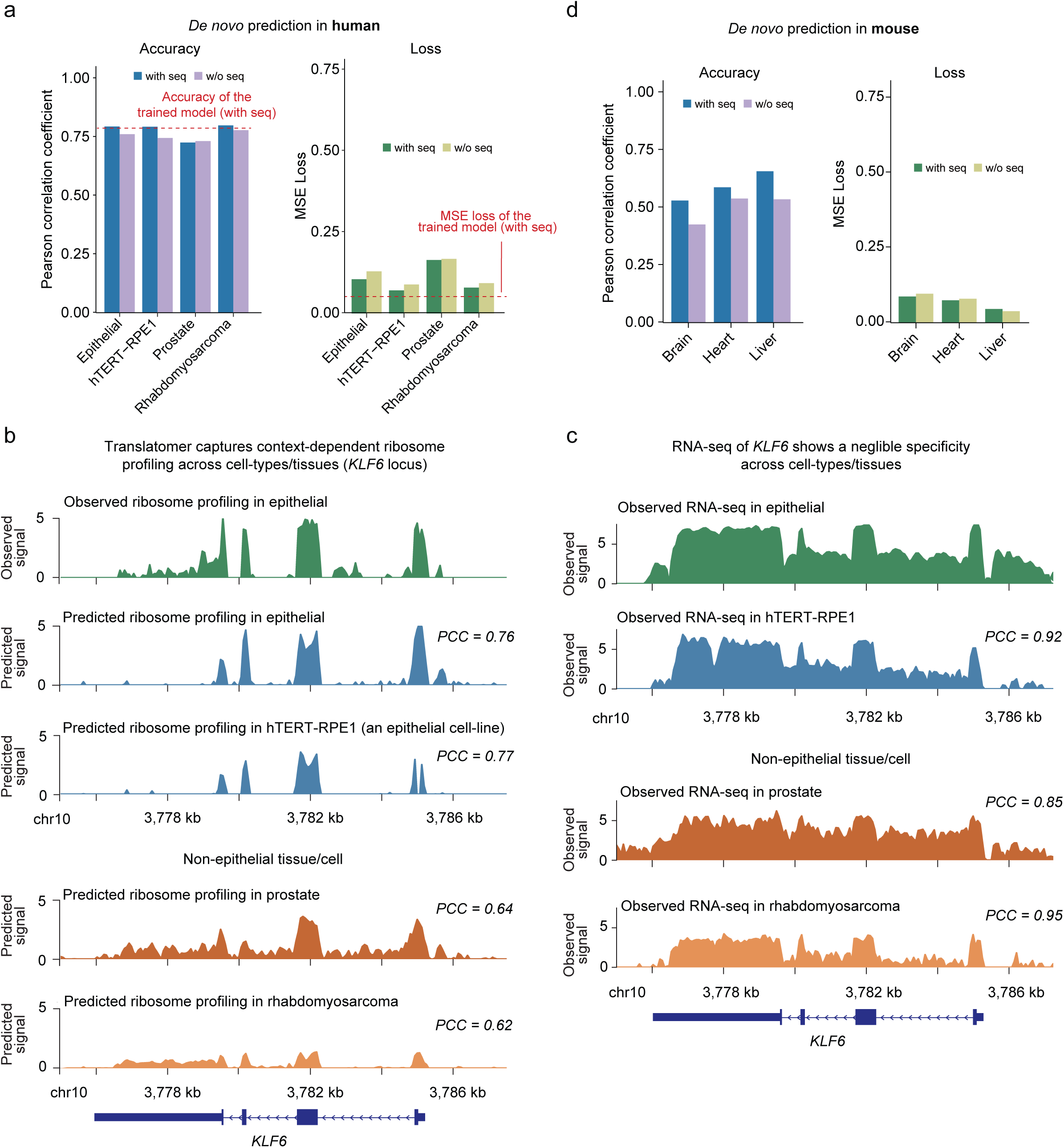
Translatomer enables accurate *de novo* prediction of ribosome profiling. a) Pearson correlation coefficient (left) and MSE loss (right) of *de novo* prediction of ribosome profiling in 4 human tissues/cell lines that are not used in the training set. The accuracy of the trained full model with sequence is indicated with the red dotted line. b) Observed and predicted ribosome profiling tracks of *KLF6* in epithelial and non-epithelial cell contexts. The Pearson correlation coefficient is calculated against the observed signal in epithelial and is labeled at the top right. c) Observed RNA-seq tracks of *KLF6* in epithelial and non-epithelial cells. The Pearson correlation coefficient is calculated against the RNA-seq signal in epithelial and is labeled at the top right. d) Pearson correlation coefficient (left) and MSE loss (right) of *de novo* prediction of ribosome profiling in mouse tissues.

In addition to the human context, we evaluated the ability of our model in *de novo* prediction for mouse translation. Notably, the model trained on human data achieves an accuracy of 0.53, 0.59 and 0.65 for mouse brain, heart, and liver in predicting ribosome profiling signals (**Fig. 2d**). These results indicate the conservation of translational regulatory rules between human and mouse, and meanwhile strongly support the accuracy and generalizability of the Translatomer model in translatome prediction.

### Sequence contribution to translatome prediction

To understand the contribution of sequence in the multi-input modeling of translation, we calculated the gradient-weighted input (gradient × input) to quantify the contribution scores of gene sequence at a base-resolution (**Methods**). We first evaluated the overall contributions of the sequence surrounding the start codon and stop codon on the translation prediction. The sequence ranging from -10 to +15 bp surrounding the start codon demonstrated the most crucial contribution to translation regulation, in concordant with the fundamental role of translation initiation in ensuring translation efficiency (**Fig. 3a**). Intriguingly, the sequence at the region of -50 to -25 bp relative to the stop codon displayed the most pronounced contribution during translation termination (**Fig. 3a**). Subsequently, we quantified the overall sequence contributions across various mRNA regions. While the coding regions exhibited the highest contribution and introns the lowest, we found that 5’ UTR wields a relatively more significant influence on translation regulation compared to 3’ UTR (**Fig. 3b**), again indicating the crucial role of the translation initiation process in modulating the overall translation strength of a gene.

**Figure 3.**
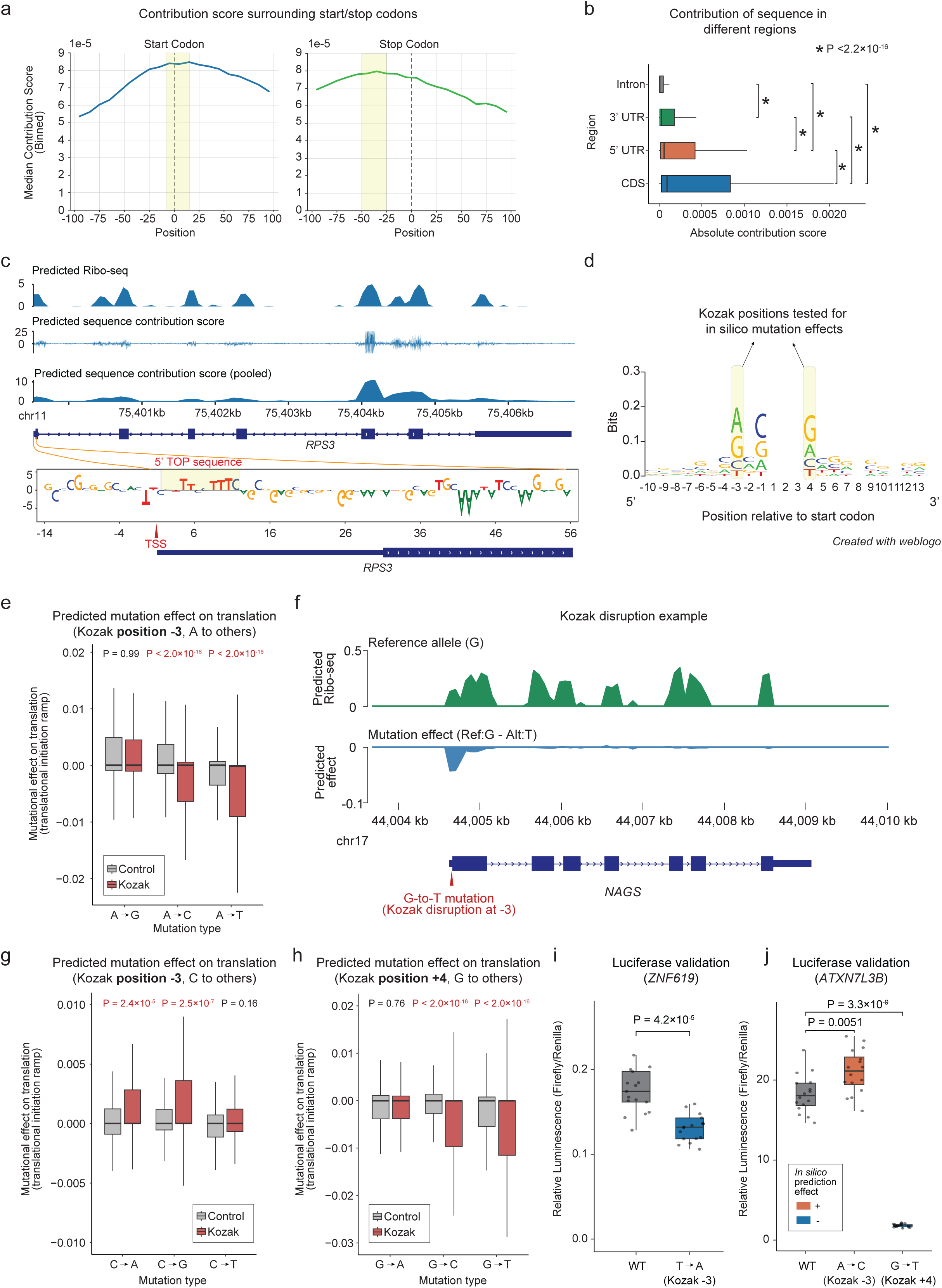
Sequence contribution on translation prediction and *in silico* mutagenesis effect estimation. a) The overall sequence contribution score surrounding start codon (left) and stop codon (right) with all the genes aggregated. The strongest contributions are highlighted in yellow. b) The absolute contribution scores of mRNA regions, including coding region, 5’ UTR, 3’ UTR, and intron. The boxplot is ordered by the median value. Pairwise p-values are calculated using Wilcoxon rank-sum test. c) Example track showing the predicted Ribo-seq signal and the sequence contribution score along the *RPS3* mRNA. The pooled sequence contribution score was calculated by aggregating the scores in bins of 128 bp. The contribution of the 5’ TOP sequence is zoomed in and visualized. d) The Kozak consensus sequence. Positions -3 and +4, which are the two most important positions for strong translation initiation, are tested for *in silico* mutagenesis effects. e) The predicted effect on translation upon the *in silico* mutagenesis from A to G (Kozak motif unaffected) and to C/T (Kozak motif disrupted) at Kozak position -3. P value was calculated using the Wilcoxon rank-sum test. f) *NAGS* gene as an example to show the *in silico* mutagenesis effect on translation of the G-to-T mutation at position -3 of the Kozak sequence. g) The predicted effect on translation upon the *in silico* mutagenesis from C to A/G (gain of Kozak motif) and to T (Kozak motif unaffected) at Kozak position -3. P value was calculated using the Wilcoxon rank-sum test. h) The predicted effect on translation upon the *in silico* mutagenesis from G to A (Kozak motif unaffected) and to C/T (Kozak motif disrupted) at Kozak position +4. P value was calculated using the Wilcoxon rank-sum test. i) Experimental verification of the mutagenesis effect based on the luciferase assay for the position -3 of Kozak sequence in *ZNF619* gene. j) Experimental verification of the mutagenesis effect based on the luciferase assay for the position -3 and +4 of Kozak sequence in *ATXN7L3B* gene. The predicted positive and negative effects on translation are denoted in red and green, respectively.

To further demonstrate the ability of the contribution score in capturing sequence elements that modulate translation efficiency, we examined the contribution score along the *RPS3* gene as an example, which is a multifunctional gene that contains the 5’ Terminal OligoPyrimidine (5’ TOP) motif^36^. The 5’ TOP motif is a cis-regulatory RNA element frequently occurring in the genes that encode proteins essential for protein synthesis, which acts as an important mechanism against environmental stress^36,37^. We show that the contribution score aligns well with the predicted ribosome footprints along the *RPS3* gene (**Fig. 3c**). We further enlarged the 5’ end of *RPS3* and visualized the contribution of each nucleotide of the first 100 bp downstream the transcription start site, and observed a strong 5’ TOP sequence consensus formed by pyrimidines (**Fig. 3c**). We also looked into *RPSA*, which is another 5’ TOP-motif-containing gene that encodes ribosomal protein, and robustly captured the relevant oligo pyrimidine sequence near the transcription start site (**Extended Data Fig. 3a**).

### *In silico* mutagenesis for predicting translational effect enabled by Translatomer and validated by luciferase reporter

To enable *in silico* perturbation on translation at a single-base resolution, we developed an *in silico* mutagenesis approach to quantify the mutational effect on the ribosome density prediction (**Methods**). We evaluated the accuracy of this Translatomer*-*based *in silico* mutagenesis assay using both computational and experimental approaches based on the Kozak sequence, which is a well-established consensus element surrounding the start codon that ensures translation initiation^38,39^. For a “strong” Kozak consensus that mediates high translation efficiency, the nucleotides at positions -3 and +4 are invariantly A/G and G, respectively^38,39^ (**Fig. 3d**).

We defined the perturbation effect as the predicted translational alteration of the start codon bin, which represents the translational initiation ramp that determines the translation efficiency of a gene^40,41^ (**Methods**). *In silico* mutations from A/G to C/T at position -3 relative to the ATG start codon led to a strong and significant negative impact on translation, while a negligible effect was observed for the counterpart of mutation located outside of Kozak sequence (**Fig. 3e-f**, **Extended Data Fig. 3b, Supplementary Table 3**). On the contrary, the mutation from C/T to A/G at the same position was predicted to increase the translation efficiency of the corresponding genes (**Fig. 3g**, **Extended Data Fig. 3c**). Additionally, we observed a dramatic reduction of translation when changing G at position +4 to other nucleotides (P= 0.76, 2 × 10^-16^ and 2 × 10^-16^ for A, C and T, respectively, **Fig. 3h**). Therefore, the predicted mutational effects on translation initiation match the expected effect of the Kozak sequence. To further estimate the mutation-induced translational effect at the level of transcript in addition to simply focusing on the translational initiation ramp, we additionally defined an *in silico* mutagenesis metric that measures the translation change of the whole coding region (**Methods**). We found a strong correlation between the predicted effects of the translation-initiation-ramp-based and the CDS-based scores (R = 0.55, P < 2.2 × 10^-16^, **Extended Data Fig. 3d**), indicating the robustness of our *in silico* mutagenesis assays in capturing translation initiation ramp and full-length coding region. Consistently, we observed highly reproducible translational effects upon the *in silico* mutagenesis at positions -3 and +4 relative to the ATG start codon, which provides further support to the accuracy of our prediction (**Extended Data Fig. 3e-f**).

Notably, despite the global down-regulation of translation upon *in silico* mutagenesis, we noticed several cases where disrupting Kozak sequence leads to an unexpected translational effect (**Supplementary Table 3**). To confirm the accuracy of these *in silico* perturbation results, we randomly selected two mutations that show unexpected predicted outcomes and carried out luciferase-based experiments to investigate the *in vivo* translational effects (**Fig. 3i-j**). For a T-to-A mutation that forms a Kozak sequence (position - 3) in *ZNF619* yet is predicted to decrease the translation, the luciferase result indeed shows a down-regulation effect (**Fig. 3i**). For an A-to-C mutation that disrupts Kozak sequence (position -3) in *ATXN7L3B* yet being predicted to increase the translation, the luciferase assay supported its positive effect on the translation of *ATXN7L3B* (**Fig. 3j**). For a G-to-T mutation that disrupts the Kozak position +4 in the same gene, where our *in silico* mutagenesis suggested a translation repression prediction that is consistent with the Kozak disruption effect, the luciferase result again supported the predicted effect on *ATXN7L3B* (**Fig. 3j**).

Collectively, the translational effects of these *in silico* mutagenesis at positions -3 and +4 are mostly consistent with the known regulatory mode of the Kozak sequence^38,39^, with rare unexpected cases that are supported by experiments. This demonstrates the accuracy of Translatomer and the *in silico* perturbation assay that we established, and meanwhile indicates the regulatory information embedded in the sequence that potentially explains the discrepancy between transcription and translation.

### Evolutionary constraint of UTR regions underlying translational regulation

Different from protein-coding sequences, 5’ UTR and 3’ UTR regions are not subjected to evolutionary constraints caused by protein sequence alterations. Here, we hypothesize that the genetic variants in these untranslated regions can also undergo evolutionary constraints by modulating the translation efficiency of mRNA transcripts. To test this hypothesis, we took advantage of our model and performed unbiased *in silico* perturbation experiments across the variants curated by the Genome Aggregation Database (gnomAD), which is the largest collection of natural human genetic variations^42^. We quantified the impacts of 851,636 gnomAD variants on translation by aggregating the ribosome density alteration along the coding region for each transcript (**Methods**). We observed that the variants with higher phyloP scores, which represent a tendency of being conserved during evolution^43–45^, show significantly increased influences on translation (**Extended Data Fig. 4a**). Compared to the variants with negative phyloP score (phyloP < 0), the variants with medium conservation signals (0 < phyloP < 5) show a 1.4-fold of increase in median effect size on translation (P < 2.2 × 10^-16^, Wilcoxon test), and the variants with high conservation (phyloP >= 5) are predicted to have a 2.4-fold of impact on translation (P < 2.2 × 10^-16^, Wilcoxon test). To avoid potential confounders caused by overall difference in phyloP scores between different mRNA regions, we further partitioned the variants based on their locations (**Fig. 4a**). We observed that the variants located in the coding regions, including synonymous and nonsynonymous mutations, are predicted to cause an overall stronger translational effect compared to those in the untranslated regions. On top of these differences between mRNA regions, the observation of the positive correlation between conservation and translational impact holds true when partitioning the variants based on their locations in 5’ UTR (N = 25,839; P < 2 × 10^-16^), synonymous (N = 131,544; P < 2 × 10^-16^), nonsynonymous (N = 343,860; P < 2 × 10^-16^) and 3’ UTR (N = 74,796; P < 2 × 10^-16^) (**Fig. 4a**).

**Figure 4.**
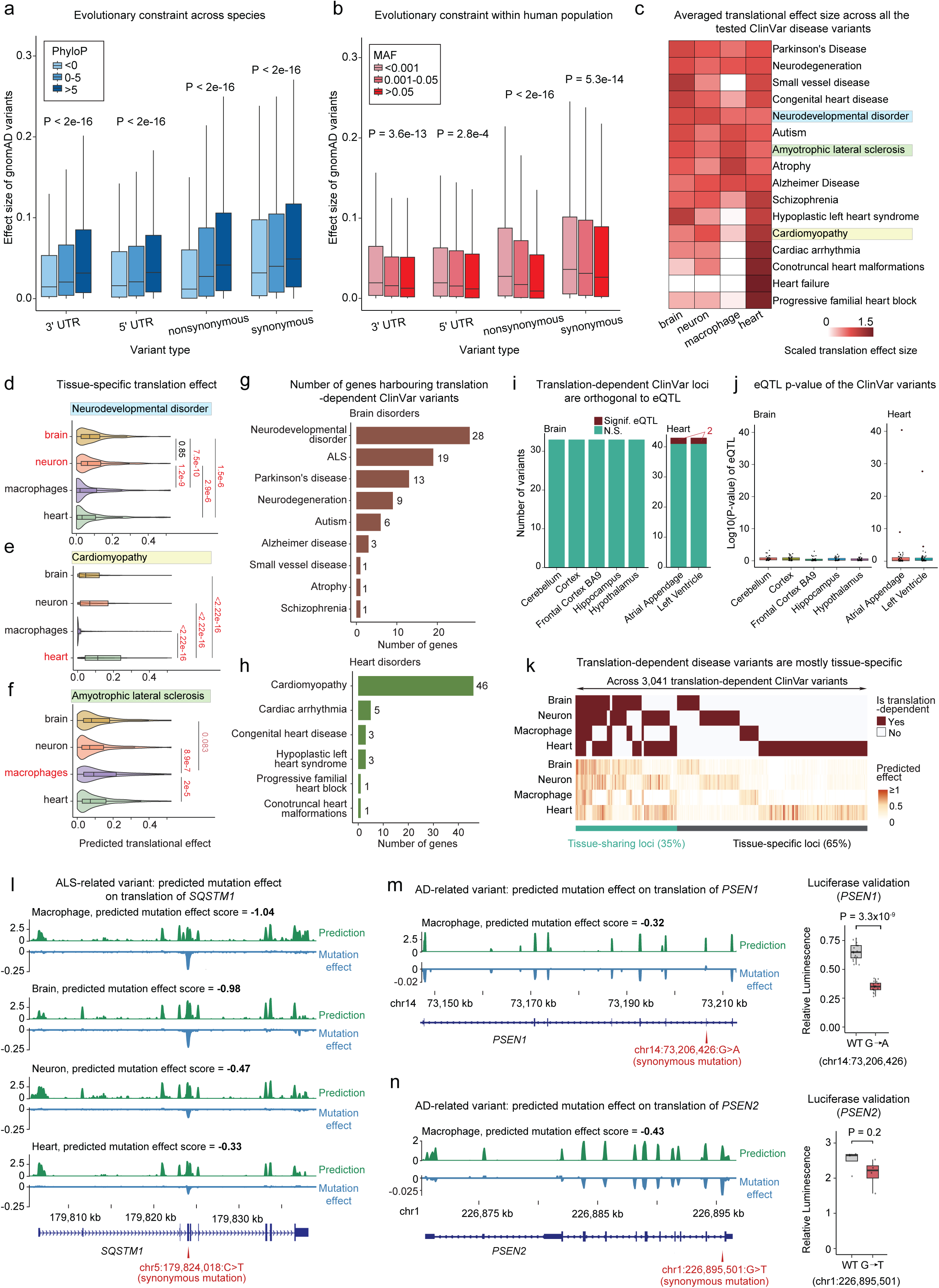
Translatomer reveals translation-dependent evolutionary constraints and interprets underpinnings of genetic diseases in a context-dependent manner. a) Effect size of *in silico* mutagenesis on translation across different ranges of phyloP score, which represents evolutionary constraint across species. The comparison is partitioned by the types of variants. P value was calculated using the Wilcoxon rank-sum test. b) Effect size of *in silico* mutagenesis on translation across different ranges of minor allele frequency, which represents evolutionary constraint within human population. The comparison is partitioned by the types of variants. P value was calculated using the Wilcoxon rank-sum test. c) The mean effect size of *in silico* mutagenesis on translation of the ClinVar variants tested for each disease in brain, neuron, macrophage and heart. d) Violin plot showing the translational effect size of the ClinVar variants associated with neurodevelopmental disorder in the four contexts. P value was calculated using the Wilcoxon rank-sum test. e) Violin plot showing the translational effect size of the cardiomyopathy-associated ClinVar variants in the four contexts. P value was calculated using the Wilcoxon rank-sum test. f) Violin plot showing the translational effect size of the ALS-associated ClinVar variants in the four contexts. P value was calculated using the Wilcoxon rank-sum test. g) Number of genes harbouring translation-dependent ClinVar variants predicted by Translatomer in brain-related disorders. h) Number of genes harbouring translation-dependent ClinVar variants predicted by Translatomer in heart-related disorders. i) Number of significant (red) and non-significant (grey) GTEx eQTLs for the variants overlapped with the identified translation-dependent ClinVar variants in brain (left) and heart (right) tissues. j) P-value of GTEx eQTLs for the variants overlapped with the identified translation-dependent ClinVar variants in brain (left) and heart (right) tissues. k) ClinVar variants dependent on tissue-sharing and tissue-specific effects on translation regulation. The upper panel shows the translation-dependent disease loci defined by Translatomer, and the bottom panel shows the translational effect predicted by the *in silico* mutagenesis. l) Example tracks of the predicted effect of an ALS-related variant, chr5:179,824,018:C>T, on the translation of *SQSTM1* gene in the contexts of macrophage, brain, neuron and heart. m) Example tracks of the predicted effect of an AD-related variant, chr14:73,206,426:G>A, on the translation of *PSEN1* gene in the contexts of macrophage (left), and the corresponding validation based on luciferase experiments (right). n) Example tracks of the predicted effect of an AD-related variant, chr1:226,895,501:G>T, on the translation of *PSEN2* gene in the contexts of macrophage (left), and the corresponding validation based on luciferase experiments (right).

Further, we interrogated the relationship between the influence of a variant on translation and its minor allele frequency (MAF), which represents the evolutionary constraint within the human population. Studies have revealed the correlation between purifying selection and MAF for genetic variations meditated by their effects on gene expression and RBP binding^46,47^. Notably, we also observed significantly elevated effects on translation for the variants with lower MAFs (**Extended Data Fig. 4b**). While the common variants (MAF > 5%) show a negligible median translational effect size (0.013), the rare variants (MAF < 0.1%) induce a considerable impact on translation (1.7 fold, P < 2.2 × 10^-16^). Moreover, the negative correlation between MAF and translational effect is consistently observed for the variants across different mRNA regions (**Fig. 4b**). Collectively, we used Translatomer to establish the role of translational regulation in driving evolution constraint, both across species and within human population as indicated by phyloP score and allele frequency, respectively.

### Cell-type/tissue-specific *in silico* perturbation identifies genetic variants relevant to complex brain and heart diseases

Having demonstrated the ability of Translatomer in capturing functionally-important sequences that are evolutionary constraint, we next sought to utilize the model to gain mechanistic insights into disease variants. We leveraged Translatomer to interrogate the disease variants located in the untranslated regions, including 5’ UTR and 3’ UTR, as well as the synonymous mutations in the coding regions. These variants do not change the sequence of protein, and the mechanisms through which they contribute to human genetic diseases remain largely elusive. We carried out *in silico* perturbation to investigate the potential mechanism mediated by translational effect for the disease-related genetic variants curated by ClinVar^48^ (**Extended Data Fig. 4c**).

Taking advantage of the ability of Translatomer in retaining cell-type/tissue-specific regulatory information, we modeled the translatome of human brain, neuron, macrophage and cardiomyocytes, and interrogated the risk variants relevant to 9 brain-related and 7 heart-related diseases (**Methods, Supplementary Table 4**). At the global level, these disease-relevant variants show context-dependent effects on the translation of the corresponding genes, as represented by the averaged effect sizes predicted by the *in silico* mutagenesis assay (**Fig. 4c**). Specifically, the variants related to brain disorders, including Parkinson’s disease (PD), neurodevelopmental disorder, and brain small vessel disease are predicted to induce an overall higher impact on the translation efficiency of the host genes in brain and neuron (**Fig. 4c-d**), while the variants in heart diseases, including cardiomyopathy, conotruncal heart malformations and heart failure, lead to the strongest translational influence in heart tissue (**Fig. 4c,e**). Notably, the translation-dependent genetic heritability of Alzheimer’s disease (AD), amyotrophic lateral sclerosis (ALS) and atrophy are primarily related to macrophage, indicating the immune component of these brain diseases (**Fig. 4c,f**).

We defined 3,041 brain- and heart-disease-relevant ClinVar variants that are predicted to impact translation efficiency of 140 genes (**Fig. 4g-h**, **Extended Data Fig. 4c-e, Supplementary Table 4),** using the gnomAD SNPs as background (**Extended Data Fig. 4c,f; Methods**). The variants predicted to affect translation account for roughly 10% to 40% among the tested ClinVar loci across the 16 brain- and heart-related diseases (**Extended Data Fig. 4g**). Of note, we excluded the nonsynonymous variants to avoid the potential confusion of mechanisms mediated by protein sequence disruption (**Extended Data Fig. 4c**). For brain disorders, we identified 341 variants in ALS, 27 in AD, 113 for PD and 40 for autism whose molecular mechanisms are likely attributed to influencing translation efficiency (**Extended Data Fig. 4d**). For heart diseases, we uncovered 58 variants for congenital heart disease (CHD), 2,236 for cardiomyopathy, 95 for cardiac arrhythmia and 33 for hypoplastic left heart syndrome that may lead to phenotypic dysfunctions via a translation-dependent mechanism (**Extended Data Fig. 4e**).

To further confirm that the mechanisms of these disease variants rely on their impacts on translation rather than transcription, we asked whether these variants show eQTL effects based on the GTEx V8 datasets^10^. For the 33 brain-disease-related variants that were tested in the GTEx dataset, none of them was identified as a significant eQTL in the 5 brain tissues (**Fig. 4i-j**). For the 43 heart-disease-related variants that were tested by GTEx, only 2 (4.65%) were identified as significant eQTLs in heart (**Fig. 4i**), with the vast majority showing negligible eQTL effects (**Fig. 4j**). Notably, we further compared the 2 significant eQTLs to their lead variants, and observed that their *p*-values were considerably lower, indicating that these 2 variants are unlikely to play causal eQTL roles (**Extended Data Fig. 4h**). These observations together strongly approve that the variants we identified are dependent on a mechanism of regulation at the translation instead of the transcription level.

Of the 3,041 disease variants identified, 1,058 (35%) affect translation efficiency in at least two tissues or cell types, while 1,983 (65%) function in a context-specific manner (**Fig. 4k, Extended Data Fig. 4i**). For instance, the ALS-related chr5:179,824,018:C>T mutation in *SQSTM1*, which encodes an autophagy receptor and has been reported to be involved in frontotemporal lobar degeneration (FTLD), AD and ALS^49–52^, leads to consistent translation decrease in macrophage, brain, neuron, and heart contexts (**Fig. 4l**). Of note, the predicted effects in macrophage (-1.04) and brain (-1.98) are over two folds stronger than the effects in the neuron (-0.47) and heart (-0.33), reflecting a context-dependent regulation for the expression of *SQSTM1* underlying ALS (**Fig. 4l**). The cardiomyopathy-related ClinVar locus, chr1:156,134,495:G>T, which is located in *LMNA* that encodes lamin A/C protein and plays a fundamental role in maintaining chromatin structure^53,54^, also shows considerable influence on the *LMNA* translation (**Extended Data Fig. 4j**). This cardiomyopathy-related variant causes the strongest translational impact in the heart context (0.98) over the other tissues/cell-types (0.24-0.42) (**Extended Data Fig. 4j**). To further confirm the reliability of the *in silico* mutagenesis in predicting the effect of disease variants, we performed luciferase experiments and verified the translational disturbance of AD-associated variants in *PSEN1* and *PSEN2* genes, respectively (**Fig. 4m-n**).

Collectively, the combination of Translatomer and *in silico* translation perturbation enables a context-dependent interrogation of disease-relevant variants, with which we provided mechanistic insights for multiple brain and heart diseases.

## Discussion

In this study, we developed a multimodal and multi-input deep learning framework, namely Translatomer, that predicts ribosome profiling signature using RNA-seq data and genomic sequence as input. We demonstrated that the regulatory information embedded in the sequence can be learned and utilized by the model to better estimate translatome from transcriptome. Combining Translatomer and the further developed *in silico* mutagenesis tool, we uncovered evolutionary constraints and disease risk variants that are ascribed to translational regulation.

Transcription and translation are two fundamental components of Central Dogma. However, the discrepancy between mRNA and protein levels has been increasingly realized^1–4^. While protein is arguably more closely related to individual phenotypes, mechanistic interpretation of disease-related variants has been largely centered on mRNA levels that are measured from RNA-seq data. For example, the number of expression QTL (eQTL) datasets (N = 933) is over 20 folds of the numbers of protein QTL (pQTL) and ribosome profiling QTL (riboQTL) datasets (N = 31 and 1, respectively), according to the Release 2.2 of QTLbase^12^. The relatively fewer research at the translation layer may partially account for the “missing regulation” of genetic diseases. Our model enables an accurate prediction of translation relying on the commonly-accessible RNA-seq data, which will facilitate the mechanistic interrogation of translation in human diseases. Moreover, mining the regulatory information embedded in the sequence may help explain non-coding disease loci and shed light on the “missing regulation” problem.

Natural language processing (NLP) models have been adapted to study the underlying regulation and context of genomic sequence, considering the intrinsic similarity between natural language and genomic sequence. The translation process in a cell is essentially a process of decoding mRNA sequence, which involves not only the sequence of coding regions, but also the regulatory information stemming from untranslated regions. Although there have been several deep learning frameworks developed for modeling ribosome profiling, our model for the first time integrates information from two modalities, including RNA-seq data and genomic sequence, to improve the accuracy of prediction. Importantly, the multi-input design of our model provides an opportunity to understand the gap between translation and transcription at the regulatory layer of sequence.

With the *in silico* mutagenesis assay developed to estimate the effect of genetic variants on translation, we systematically investigated the relationship between evolutionary constraint and translational effect. Although UTR regions do not immediately alter protein sequence, there are various regulatory elements in UTRs that modulate the translation efficiency of the host genes. We examined two relevant but different aspects, including phyloP and MAF, which represent evolution signals at the levels of inter-species and intra-human populations, respectively. For both results, our model revealed that the effect on translation mediates the evolutionary constraints of the variants in the UTR regions.

Importantly, our model sheds light on 3,041 non-coding variants that are reported to be relevant to 16 diseases, including ALS, schizophrenia, autism, AD, small vessel disease, cardiomyopathy, congenital heart disease and more. We also identified 2,899 synonymous variants, which do not alter protein sequence, that contribute to human diseases by influencing the translation efficiency of the coding regions. We excluded the possibility that these variants function through transcriptional effect based on our observation that they are eQTL-independent. We show that the risk variants of a certain disease tend to show an overall stronger effect on translation in the relevant tissue. For example, the translatome of neuronal diseases, including ALS, schizophrenia and autism, tends to be more responsive to the genetic alterations in brain and neuron cells, while small vessel disease, heart failure and cardiomyopathy are more relevant to heart. Interestingly, AD-related variants are predicted to cause stronger translational alteration in macrophage, instead of in neuron or brain, in line with the recent realization that AD is an immune onset disorder^55^. Therefore, our results provided mechanistic insights into the disease variants that affect translation in a context-specific fashion.

Overall, Translatomer provides not only a framework for *de novo* prediction of translatome from the easily-accessible RNA-seq data, but also an interface for the interrogation of the effect of genomic sequence on translation process. We envision that our model will help accelerate our understanding of functional genomics and missing regulation in the field of disease genetics.

## Methods

### Processing of ribosome profiling and the paired RNA-seq data

The ribosome profiling data and RNA-seq data used in this study were obtained from publicly available sources on the Gene Expression Omnibus (GEO) website (www.ncbi.nlm.nih.gov/geo). In total, we curated 40 paired datasets in raw FASTQ format (**Supplementary Table 1, Supplementary Table 2**), comprising 37 human datasets and 3 mouse datasets.

For the ribosome profiling data preprocessing, we initiated the pipeline by excluding rRNA and tRNA sequences. Reference rRNA and tRNA sequences were downloaded from the UCSC table browser. We employed Bowtie2^56^ (v.2.2.5) to align the raw Ribo-seq data against the reference rRNA and tRNA sequences. Subsequently, the resulting reads that did not align to rRNA and tRNA were mapped to the hg38 human reference genome and the mm10 mouse genome, respectively, utilizing Bowtie2 with the “--local” parameter. We employed SAMtools^57^ (v.1.6) to convert the mapped reads into BAM (Binary Alignment Map) format, followed by sorting the resulting BAM files. To ensure signal consistency across diverse cell lines, we adopted a strategy of merging FASTQ data from various replicates. Finally, the processed BAM file was transformed into the bigwig format employing bamCoverage^58^ (v.3.5.1), with the “--binSize 1” parameter.

For the RNA-seq data preprocessing, quality control was first performed using FASTP^59^ (v.0.23.2), and low-quality reads were trimmed or removed. Subsequently, we performed sequence alignment using Bowtie2 with the command “--local” to align the processed RNA-seq reads to the reference genome. The aligned reads were then converted into BAM format using SAMtools. Then we merged the BAM files generated from different replicates into a single, unified BAM file. Finally, we utilized the bamCoverage tool with the “--binSize 1” parameter to create bigwig files.

### Input and output preprocessing for the model

#### Input

We defined gene regions as the target regions for each input. Specifically, we extracted the genomic intervals including chromosomes, start site, and end site from the gene annotation file in the GFF format downloaded from GENCODE (v43) by selecting ’protein_coding’ in the column ’gene_type’, and subsequently saved into a BED file, serving as the target regions of our model. To define the region for training, we fed in the start and end positions and obtained a region of 65,536 bp centered at the middle of each gene. If a gene is shorter than 65,536 bp, the region upstream of TSS and downstream of TES will also be included.

For the processing of gene sequence input, we used the reference genome sequence (hg38 and mm10) from the UCSC Genome Browser database. The original FASTA file includes four types of nucleotides and ‘N’ for unknown nucleotides. We retained the ‘N’ nucleotides and encoded them as the fifth ‘nucleotide’ type. We employed one-hot encoding to represent DNA sequences, where each nucleotide was mapped to a 5-dimensional vector (A = [1, 0, 0, 0, 0], T = [0, 1, 0, 0, 0], C = [0, 0, 1, 0, 0], G = [0, 0, 0, 1, 0], N = [0, 0, 0, 0, 1]).

RNA-seq sequence coverage values of target regions are extracted using the Python package pyBigWig from the bigwig files and resized to 65,536 bp utilizing the *resize* function within the *Interval* module of the kipoiseq Python package. The same procedures were used to obtain aggregated RNA-seq signals and were subsequently used as input for the model. To mitigate the effect of extreme values, we performed a log(x + 1) transformation on both RNA-seq and Ribo-seq bigWig files.

The input consists of RNA-seq and gene sequence. We combined the one-dimensional RNA-seq value and five-dimensional gene sequence, which resulted in a six-dimension vector (N × 65,536 × 6). The combined features were transposed into dimension (N × 6 × 65,536) as inputs for the Translatomer model, where N is the total sample size.

#### Output

Following the same way of preprocessing for RNA-seq data, we extracted sequencing coverage values of Ribo-seq from bigwig files, which were initially of length 65,536. Then the 65,536 dimensions were aggregated into values of length 1,024 with a bin size of 64. These aggregated Ribo-seq values with shape (N × 1024) were set as the output for the Translatomer model.

### Model architecture of Translatomer

The Translatomer model takes the gene sequence and RNA-seq as inputs, and Ribo-seq as output. The Translatomer architecture consists of an input layer, a transformer tower, and an output layer^27^. The architecture was implemented in Pytorch.

The input layer is composed of a one-layer one-dimension convolution block (nn.Conv1d with kernel size = 129, stride = 128, and padding = 64), a Batch Normalization layer, and a ReLU activation layer. The input layer transforms the input dimension from (N × 6 × 65,536) to (N × 512 × 512) to feed into the transformer tower, where N is the batch size.

The transformer tower is an attention module composed of a 12-layer Transformer EncoderLayer block (nn.TransformerEncoderLayer with hidden dimension is 512 and number of attention heads is 8). Relative key-query positional embedding is also applied and the maximum length is set to be 512. The Transformer tower is responsible for extracting the features from input RNA-seq and DNA sequences. The dropout rate is set to 0.1 in the positional encoding and within each Transformer layer.

The output layer is composed of a one-layer one-dimension convolution block (nn.Conv1d with kernel size = 3, stride = 1, and padding = 1) and a fully connected layer (nn.Linear), and a Dropout layer with dropout rate of 0.1, ReLU activation. The output layer transforms the features extracted by the transformer tower into output (N × 1 × 1024) and squeezes into (N × 1024) as predicted ribosome profiling coverage.

The mean-squared error (MSE) loss is calculated between the predicted and the experimental Ribo-seq tracks (ground truth). The average Pearson correlation coefficient (PCC) is calculated to evaluate the model.

### Model training, validation and testing

The training data comprises the reference gene sequence and RNA-seq signal as inputs, and Ribo-seq signal output, where the pairwise RNA-seq and Ribo-seq were obtained from 33 human tissues and cell lines. Genes located in chromosomes 10 and 15 were used as the validation set and the test set, respectively, and the genes in the remaining chromosomes were utilized for model training.

The Translatomer model was trained within Pytorch Lightning (v.1.7.6) framework with a batch size of 32, AdamW optimizer (learning rate = 1e-5), a weight decay of 0.05, and early stopping with a patience of 8 epochs. A cosine learning rate scheduler with 2,000 warm-up steps was employed for stabilizing training. Two types of data augmentations were applied, including reverse complementation of the sequence and flipping of the target Ribo-seq and RNA-seq values. The model achieved minimal validation loss after 38 epochs of training on a single NVIDIA Tesla 80G A100 GPU.

### Benchmark with the state-of-the-art methods

We compared our model with two state-of-the-art methods for modeling ribosome density: iXnos^26^, a machine learning based approach, and RiboMIMO^31^, a deep learning architecture. We used ribosome profiling data from K562, epithelial cells, brain, and HepG2 human cell lines for the model comparison. Our model takes both RNA-seq and full mRNA sequence (5’ UTR, coding region, and 3’ UTR) as input, while the other two models take the coding region of mRNA sequence as input. We strictly followed the same mRNA sequence encoding methods as the original papers for iXnos and RiboMIMO, with both methods taking a preprocessing step to encode the coding sequence as 3-bp codons. In this scheme, codons are represented as one-hot vectors of 64 dimensions, while nucleotides are formed by concatenating three one-hot vectors representing the four nucleotide types (A, C, G, and T). Similarly, amino acids and stop codons are represented as one-hot vectors of 21 dimensions. To robustly evaluate the performance of the models, we used an 11-fold cross-validation by holding out genes in different chromosomes as validation and testing datasets at each fold.

For a fair and straightforward comparison under a consistent feature condition, we adjusted the input and output sequence lengths for each of the models. We utilized an input length of 5,120 for our model, and 1,024 as the length of the output Ribo-seq signal. For the other codon-based models, both the input and output lengths were set to 1,707, corresponding to a 5,121 sequence length (1,707 x 3 bp). This adjusted data size allows for a more focused comparison of model performance. The iXnos and RiboMIMO models were implemented based on the publicly available code provided from the original papers. We used the best hyperparameter settings as mentioned in the papers for each of the models, and then evaluated model performance using the average Pearson correlation coefficient (PCC). The comparison gave rise to accuracies ranging from 0.31 to 0.49 for iXnos, and 0.45 to 0.55 for RiboMIMO, which were both comparable to the accuracies reported by the original papers^26,31^.

### Contribution score of gene sequence

We employed a backpropagation algorithm to compute the gradient of the model’s output in correspondence to the input sequence. Specifically, we used the gradient × input^60^ approach to compute the contribution score of each position in the input sequence of the model. To remove the effect of RNA-seq input, we normalized the input × gradient results over RNA-seq.

### *In silico* mutagenesis effect scoring

For each variant and locus, we calculated an *in silico* mutagenesis effect score to estimate its impact on translation. This score is defined as the difference between the prediction generated from the original input (reference allele) and the prediction derived from the perturbed input (alternative allele). The representation is expressed as follows:

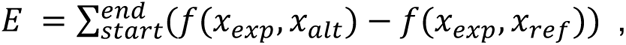

where *x_exp_* represents the RNA-seq signals, *x_alt_* represents the modified sequence with the alternative allele, while *x_ref_* represents the reference genome sequence. The [start:end] notations denote the start and end bins encompassing the start codon bin and all coding sequence (CDS) regions, respectively.

### Mutational effect evaluation against Kozak motifs

We first utilized the Kozak motif, which is a well-known sequence consensus that drives translation initiation, as a positive control to evaluate the performance of the *in silico* mutagenesis tool that we established. For each gene in the genome, the position -3 relative to the ATG start codon, which is crucial in the Kozak motif, was mutated to the other three nucleotides, and the mutation effects were calculated. For example, for a gene with an “A” at position -3, the *in silico* mutation effect upon *in silico* mutations to G, T, and C were quantified, respectively. Additionally, for all the genes with a “G” at their positions +4, we performed *in silico* mutation to A, T, and C, and estimated their effects on the translation predictions. The general effects of *in silico* mutations at these two positions were examined across all the genes in the genome. We tried two different regions to define *in silico* mutagenesis effect on translation, one was based on the start codon bin that represents the translation initiation ramp, and the other one was based on the whole coding region. The mutational effects based on these two regions were highly consistent with each other, and were both in concordance with the known regulatory roles of the Kozak sequence.

### Evolutionary constraints assessed by *in silico* mutagenesis effect on translation

We tested the potential relationship between the effects on translation efficiency and the evolutionary constraints for the variants collected by the Genome Aggregation Database (gnomAD)^42^. This analysis was confined to the genetic variants located in the gene regions, including 5’ UTRs, coding regions, and 3’ UTRs, which resulted in a total of 851,636 variants. Evolutionary constraint was represented by minor allele frequency (MAF), which was obtained from the gnomAD annotation, as well as phyloP score, which was downloaded from the UCSC genome browser (basewise conservation scores of 99 vertebrate genomes with human). MAF and phyloP represent the evolutionary constraint within human populations and across species, respectively. For each of these variants, we calculated the effect on translation based on the *in silico* tool detailed above, and then performed association analysis with MAF and phyloP, respectively. The comparison was further partitioned based on the mRNA location of variants, including 5’ UTRs, 3’ UTRs, and the synonymous and nonsynonymous variants in coding regions.

### Experimental validation using luciferase reporter assays

To examine the effect of mutations on translation, we constructed reporter plasmids based on pmirGLO Dual-Luciferase miRNA Target Expression Vector (Promega). We used ClonExpress II One Step Cloning Kit (Vazyme, C112) to insert the mRNA 5’ UTR and CDS regions that contain the mutations of interest to the upstream of FLAG-T2A tag. Subsequently, a mutation primer was designed and synthesized to introduce site-specific mutations based on the wildtype plasmids. The mutagenesis PCR was performed using the Q5® High-Fidelity 2X Master Mix, followed by DpnI digestion of the original plasmid. Gel purification and re-ligation steps were carried out. The wild type and mutation plasmids were then purified using the EasyPure® HiPure Plasmid MiniPrep Kit (Transgen, EM111-01).

HEK-293T cells were cultured in DMEM medium, supplemented with 10% FBS (Gibco) and 1% glutaMAX (Gibco). Transfection of HEK-293T cells was performed using lipo3000 (Invitrogen, L3000001), following the instructions provided in the manuscript. Briefly, the plasmid and P3000 were diluted in Opti-MEM, while lipo3000 was diluted in a separate tube of Opti-MEM. The two tubes of medium were mixed, and the resulting complex was added to the cells.

After 72 hours of transfection with the reporter plasmid, the activity of firefly luciferase was analyzed using the Dual-Glo® Luciferase Assay System (Promega, E2920), along with the Renilla luciferase control, to assess the transfection efficiency and normalization.

## Acknowledgements

We thank Dr. Xiaoyu Li for sharing the luciferase reporter plasmid for the experimental validation of the identified disease risk loci. We thank Dr. Lei Hou for the discussion and suggestions. We thank the members in the Xiong Lab for discussion and suggestions throughout the project. We thank the support from the core facilities and computing platform of Liangzhu Laboratory at Zhejiang University. This work was supported by the National Natural Science Foundation of China nos. 32370609 and 92353301 to X.X. and no. 82303974 to J.L., and the funding from Liangzhu Laboratory at Zhejiang University and the State Key Laboratory of Transvascular Implantation Devices.

## Author Contributions

This study was designed by J.H., L.X., and X.X., and directed and coordinated by X.X.. J.H., C.L. trained and fine-tuned the model with the help and under supervision of J.N., K.D., L.X., Y.M., M.K. and X.X.. S.S. and K.C. performed experimental validation under supervision of J.L. and X.H.. All authors participated in the discussion of the project. J.H., L.X. and X.X. wrote the manuscript.

## Competing interests

The authors declare no competing interests.

## Code availability

The code will be available upon formal publication.

## Data availability

All data is publicly available at GEO, with detailed information and accession numbers available in Table S1 and Table S2.

## Figure legends

**Extended Data Figure 1.**
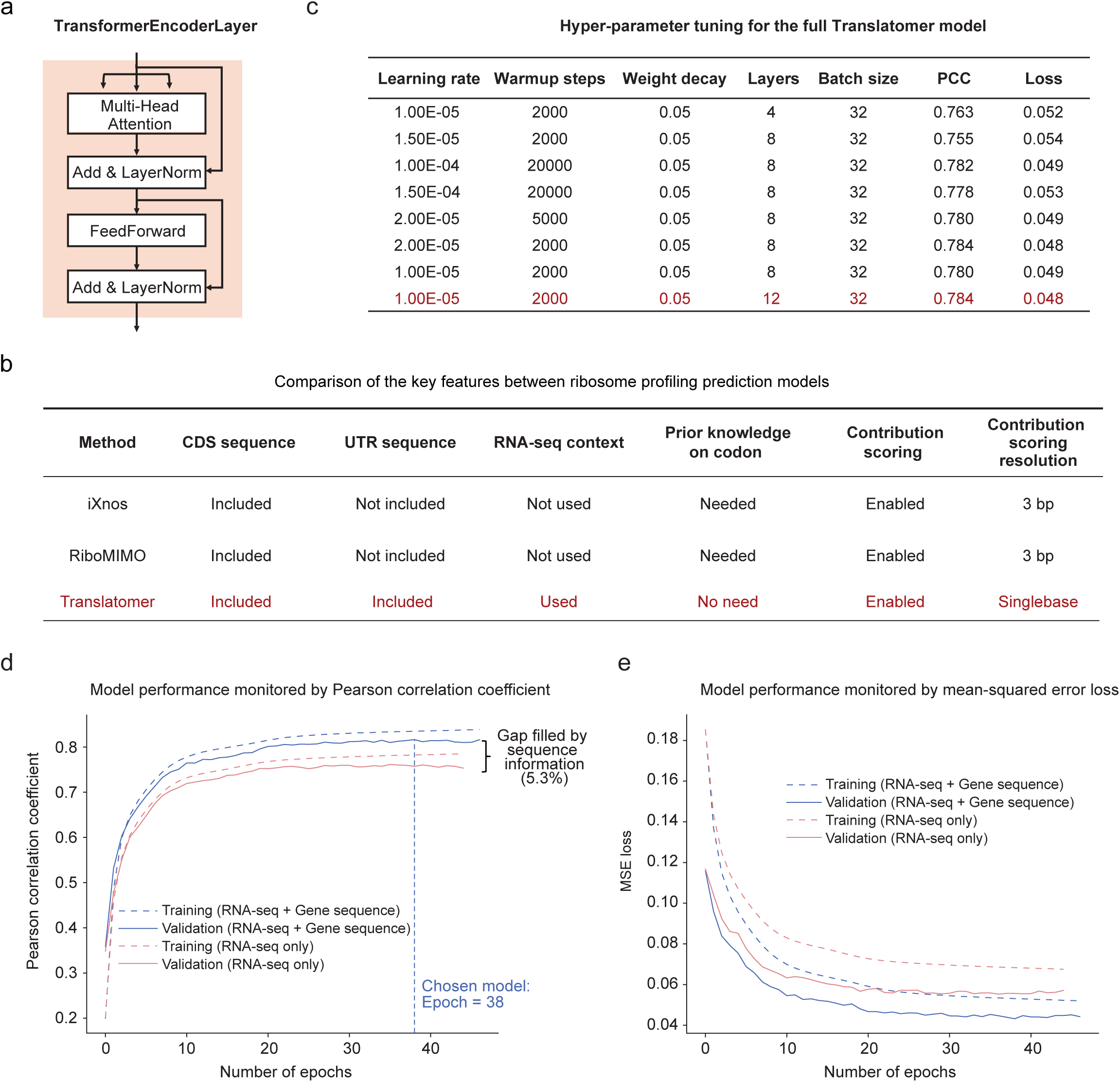
Features and performance of Translatomer model. a) Sketch plot showing the architecture of the transformer layer used in this study. The full Translatomer model is shown in Figure 1a. b) Comparison of the key features between Translatomer and other ribosome profiling prediction models. c) Table showing the performance of different hyper-parameters tested during model construction using the datasets from 33 tissues and cell lines. d) Pearson correlation coefficient (PCC) increases and converges as the number of training epoch increases. Multi-input and single-input models are in blue and red, respectively. Training accuracy is represented by a dotted line and validation accuracy is represented by a solid line. The accuracy difference between multi-input and single-input models is calculated based on the validation accuracy. e) Mean-squared error loss decreases and converges upon the increase of training epochs.

**Extended Data Figure 2.**
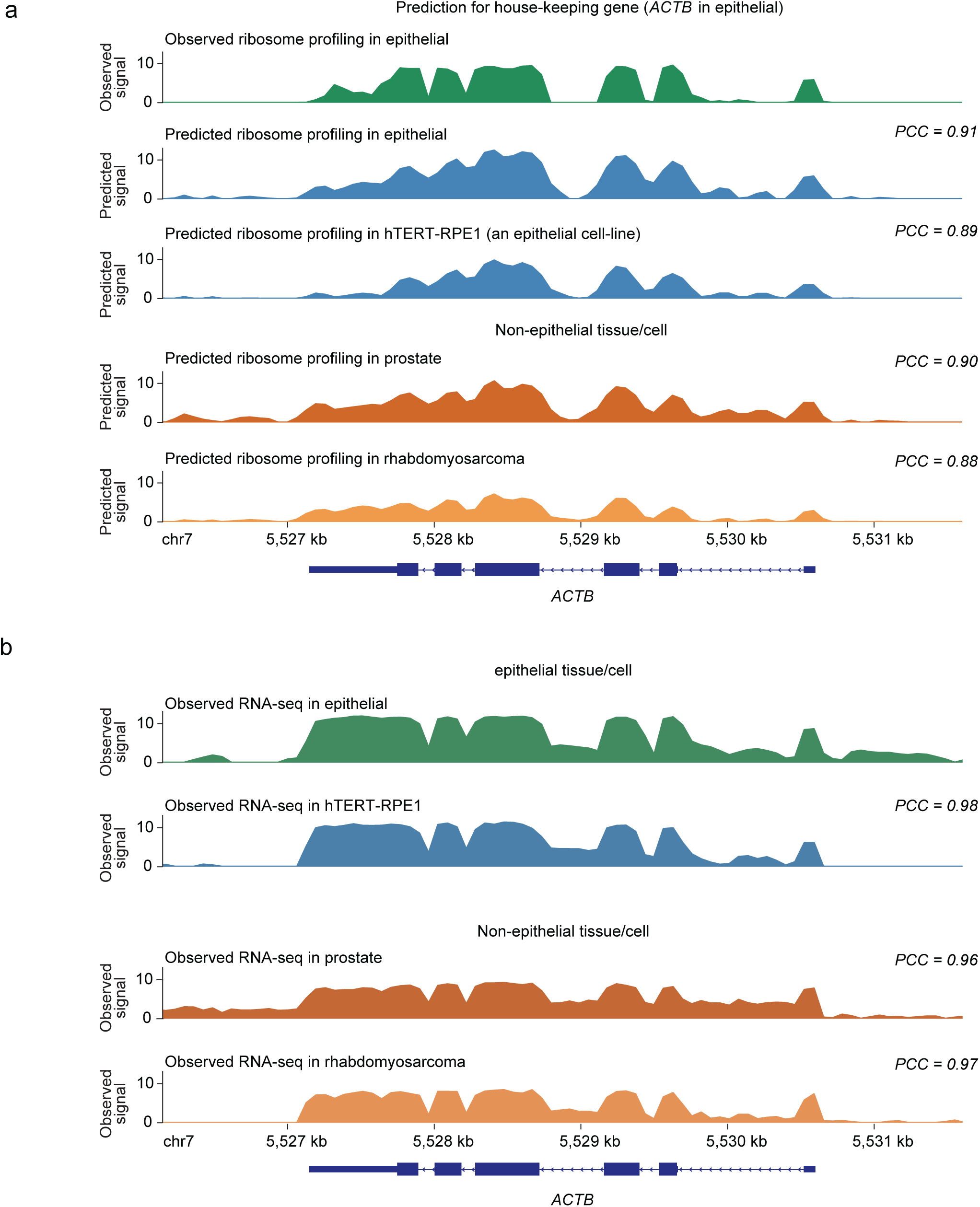
Translatomer accurately predicts ribosome profiling signal for new data. a) Observed and predicted ribosome profiling tracks in epithelial cells and non-epithelial cells for the *ACTB* gene. The Pearson correlation coefficient against the observed ribosome profiling in epithelial is labeled at the top right. b) Observed RNA-seq tracks of *ACTB* in epithelial and non-epithelial cells. The Pearson correlation coefficient is calculated against the RNA-seq signal in epithelial and is labeled at the top right.

**Extended Data Figure 3.**
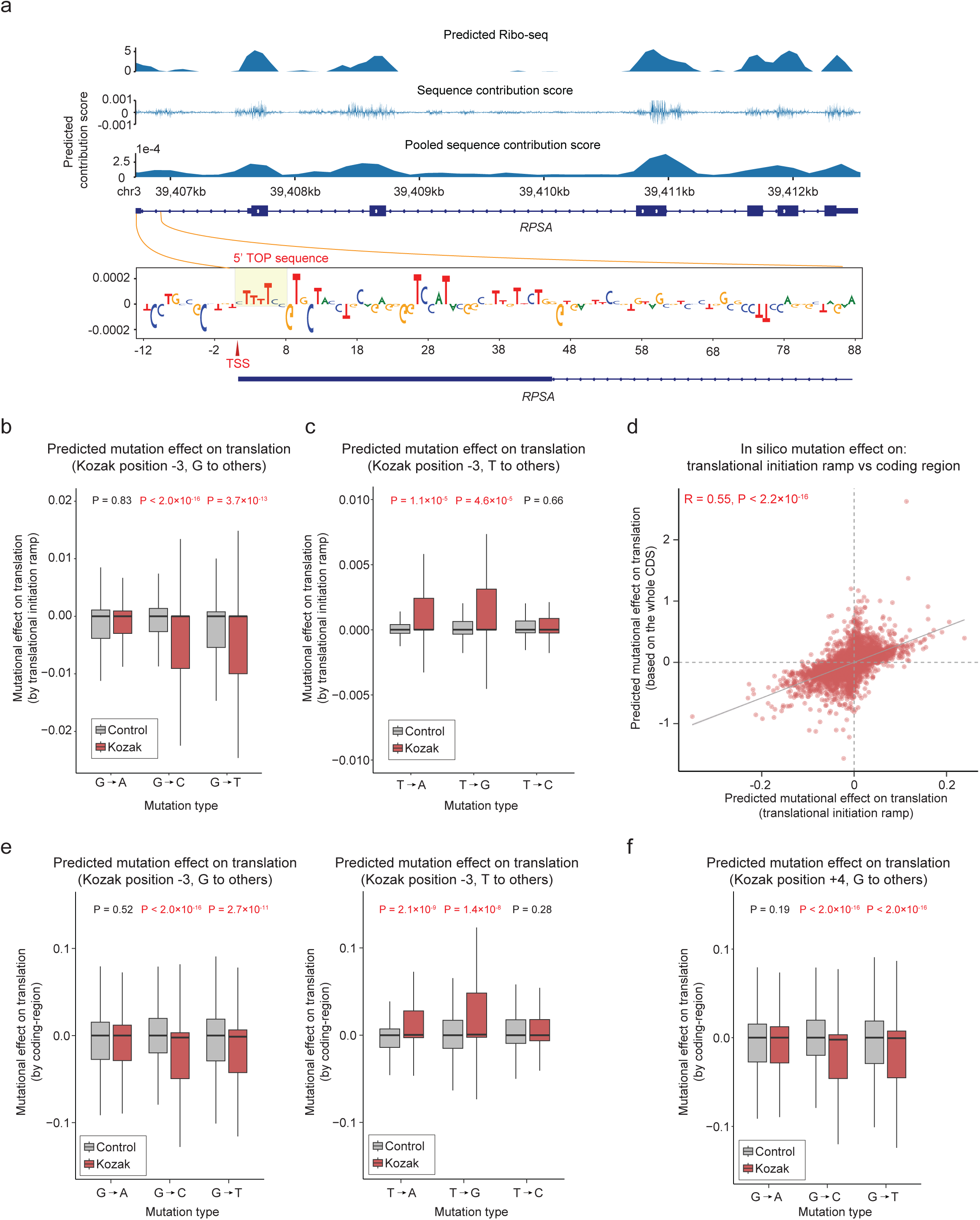
Validation of Translatomer based on *in silico* mutagenesis of Kozak sequence. a) Example track showing the predicted Ribo-seq signal and the sequence contribution score along the *RPSA* mRNA. The pooled sequence contribution score was calculated by aggregating the scores in bins of 128 bp. The contribution of the 5’ TOP sequence is zoomed in and visualized. b) The predicted effect on translation upon the *in silico* mutagenesis from G to other nucleotides at position -3. P value is calculated using Wilcoxon rank-sum test. c) The predicted effect on translation upon the *in silico* mutagenesis from T to other nucleotides at position -3. d) Scatter plot showing the correlation between the *in silico* mutagenesis effects based on the translation initiation ramp (x-axis) versus the whole coding region (y-axis). The R and p-value of the correlation analysis were shown. e) The predicted effect on translation upon the *in silico* mutagenesis from G (left) and T (right) to other nucleotides at position -3. The effect was estimated based on the whole coding region. f) The predicted effect on translation upon the *in silico* mutagenesis from G to other nucleotides at position +4. The effect was estimated based on the whole coding region.

**Extended Data Figure 4.**
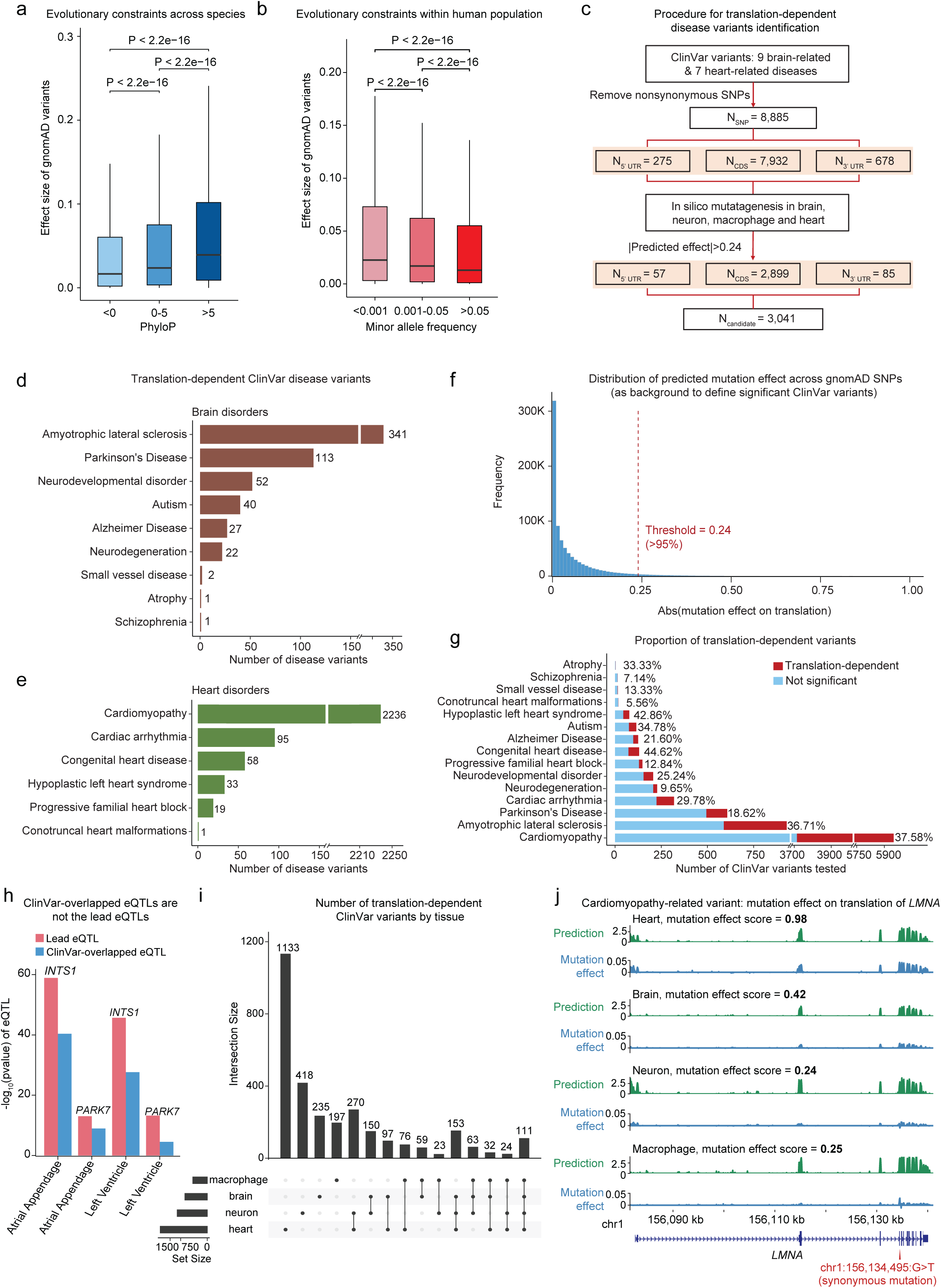
Evolutionary constraints interrogation and disease variants interpretation by Translatomer. a) Effect size of *in silico* mutagenesis on translation across different ranges of PhyloP score, which represents evolutionary constraint across species. P value was calculated using the Wilcoxon rank-sum test. b) Effect size of *in silico* mutagenesis on translation across different ranges of minor allele frequency, which represents evolutionary constraint within human population. P value was calculated using the Wilcoxon rank-sum test. c) Procedure for the identification of translation-dependent ClinVar variants based on *in silico* mutagenesis. d) Number of translation-dependent ClinVar variants identified by Translatomer in brain-related disorders. e) Number of translation-dependent ClinVar variants identified by Translatomer in heart-related disorders. f) Distribution of the absolute *in silico* mutagenesis effect on translation across the gnomAD variants. A threshold of 0.24, which corresponds to the effect ranking at the top 5%, is selected to define the candidate variants that influence translation efficiency. g) Number of ClinVar variants that are dependent (red) and independent (blue) of their impacts on translation. The percentage of the translation-dependent variants for each disease is labeled. h) The translation-dependent ClinVar variants showing eQTL significance are not lead eQTL SNPs in the corresponding loci. i) The number of translation-mediated variants identified for each disease curated by the ClinVar database. The sharing of the cell type/tissue contexts is shown at the bottom. j) The example tracks of the chr1:156,134,495:G>T effects on the translation of *LMNA* gene in the contexts of heart, brain, neuron and macrophage.

## Notes

### Competing Interest Statement

The authors have declared no competing interest.

